# A group B streptococcal type VII secreted LXG toxin mediates interbacterial competition and colonization of the female genital tract

**DOI:** 10.1101/2024.06.10.598350

**Authors:** Alyx M. Job, Kelly S. Doran, Brady L. Spencer

**Affiliations:** Department of Immunology and Microbiology, University of Colorado Anschutz, Aurora, CO, USA

**Keywords:** *Streptococcus agalactiae*, group B *Streptococcus*, GBS, Type VII Secretion System, LXG toxins, interbacterial competition, vaginal colonization

## Abstract

Group B *Streptococcus* (GBS) asymptomatically colonizes the vagina but can opportunistically ascend to the uterus and be transmitted vertically during pregnancy, resulting in neonatal pneumonia, bacteremia and meningitis. GBS is a leading etiologic agent of neonatal infection and understanding the mechanisms by which GBS persists within the polymicrobial female genital mucosa has potential to mitigate subsequent transmission and disease. Type VIIb secretion systems (T7SSb) are encoded by Firmicutes and often mediate interbacterial competition using LXG toxins that contain conserved N-termini important for secretion and variable C-terminal toxin domains that confer diverse biochemical activities. Our recent work characterized a role for the GBS T7SSb in vaginal colonization and ascending infection but the mechanisms by which the T7SSb promotes GBS persistence in this polymicrobial niche remain unknown. Herein, we investigate the GBS T7SS in interbacterial competition and GBS niche establishment in the female genital tract. We demonstrate GBS T7SS-dependent inhibition of mucosal pathobiont *Enterococcus faecalis* both *in vitro* using predator-prey assays and *in vivo* in the murine genital tract and found that a GBS LXG protein encoded within the T7SS locus (herein named group <u>B</u> streptococcal <u>L</u>XG <u>T</u>oxin <u>A</u>) that contributes to these phenotypes. We identify BltA as a T7SS substrate that is toxic to *E. coli* and *S. aureus* upon induction of expression along with associated chaperones. Finally, we show that BltA and its chaperones contribute to GBS vaginal colonization. Altogether, these data reveal a role for a novel T7b-secreted toxin in GBS mucosal persistence and competition.

**Importance:** Competition between neighboring, non-kin bacteria is essential for microbial niche establishment in mucosal environments. Gram-positive bacteria encoding T7SSb have been shown to engage in competition through export of LXG-motif containing toxins, but these have not been characterized in group B *Streptococcus* (GBS), an opportunistic colonizer of the polymicrobial female genital tract. Here, we show a role for GBS T7SS in competition with mucosal pathobiont *Enterococcus faecalis*, both *in vitro* and *in vivo*. We further find that a GBS LXG protein contributing to this antagonism is exported by the T7SS and is intracellularly toxic to other bacteria; therefore, we have named this protein group <u>B</u> streptococcal <u>L</u>XG <u>T</u>oxin <u>A</u> (BltA). Finally, we show that BltA and its associated chaperones promote persistence within female genital tract tissues *in vivo.* These data reveal previously unrecognized mechanisms by which GBS may compete with other mucosal opportunistic pathogens to persist within the female genital tract.

## Introduction

Interbacterial competition is necessary for bacterial niche establishment, particularly in polymicrobial mucosal environments within the host (1). Bacteria utilize numerous systems to compete with their non-kin neighbors, including nutrient restriction, production of bacteriocins or antibiotics, as well as secretion systems that facilitate export of toxic effectors (2). The Type I, Type IV, and Type VI Secretion systems in Gram-negative bacteria have been particularly well-studied for their roles in interbacterial competition by secreted toxins (3). However, mechanisms of interbacterial antagonism by specialized secretion systems in Gram-positive bacteria have been less characterized.

Type VIIb secretion systems are encoded by Firmicutes and are comprised of core machinery proteins (membrane-bound EsaA, EssA, EssB and cytoplasmic EsaB) as well as a membrane-associated FtsK/SpoIII ATPase, EssC, which drives the export of small ɑ-helical proteins and effectors (4). WXG100 proteins (named for their central Tryptophan-X-Glycine motif and 100 amino acid structure) are canonical substrates of the T7SS and have been implicated in virulence in numerous Gram-positive species (5–7). LXG toxins are also common substrates of the T7SSb and are characterized by ɑ-helical N-terminal structures containing a Leu-X-Gly motif. These N-terminal domains have been shown to interact with adjacently encoded ɑ-helical chaperones, or <u>L</u>XG-associated <u>ɑ</u>-helical <u>p</u>roteins (Laps), and this complex is hypothesized to be important for recognition and trafficking to the secretion machinery (4, 8). The C-terminal domain of these proteins often confers toxic activity and its structure is variable between LXG proteins depending on their biochemical function (4). Indeed, LXG toxins across and within genera and species are often biochemically diverse and the while functions of many remain unknown, the few that have been studied may act as nucleases, NADases, or phospholipases (4, 9, 10). In several Gram-positive bacteria, T7SSb-secreted LXG toxins has been shown to exhibit toxicity and/or contribute to interbacterial competition (8, 9, 11–15) and T7SSb has been hypothesized to have evolved for interbacterial antagonism, promoting niche establishment by both commensals and pathogens (16). Therefore, study of these toxins across additional T7SSb-encoding opportunistic pathogens is needed to better understand mechanisms of competition in the mucosa.

*Streptococcus agalactiae* or group B *Streptococcus* (GBS) is a Gram-positive pathogen that asymptomatically colonizes the vaginal tract but causes infection upon ascension to higher genital tract tissues, such as the uterus (17). During pregnancy, GBS vaginal colonization and ascending infection can result in adverse pregnancy outcomes (including stillbirth, pre-term birth, and chorioamnionitis) as well as neonatal pneumonia, bacteremia, and meningitis upon vertical transmission (18). As vaginal colonization is a prerequisite for development of early onset GBS disease in infants (19), a better understanding of the bacterial factors that promote persistence of GBS within the vaginal mucosa is needed. While the microbiota often confers colonization resistance in mucosal niches (20, 21), GBS is not only capable of establishing persistent colonization, but is also known to perturb the vaginal microbiota during colonization (21–23). Thus, the specific mechanisms by which GBS competes with the microbiota and/or mucosal pathobionts to persist within this polymicrobial niche warrant further study.

We previously characterized the GBS T7SSb and identified four T7SS subtypes across GBS clinical isolates based on variation within the C-terminus of the EssC ATPase and unique downstream repertoires of putative effectors (24). Despite this effector diversity across subtypes, three of the four GBS T7SSb subtypes’ loci display a conserved synteny downstream of the T7SS machinery genes, encoding for two putative chaperones and a putative LXG toxin, similar to those described in other Gram-positive species (8, 25). We demonstrated that the highly prevalent subtype I GBS T7SS contributes to vaginal colonization and ascending infection, *in vivo* (7, 24). However, the mechanism by which the subtype I T7SS establishes this niche had not been determined. We hypothesize that the GBS T7SSb and, specifically, secreted LXG toxins may promote interbacterial antagonism, facilitating persistent GBS vaginal colonization and infection. Herein, we identify a role for the GBS T7SS in interbacterial competition with *Enterococcus faecalis* both *in vitro* and *in vivo* in the female genital tract. We further identify a GBS LXG protein encoded within the T7SS subtype I locus that contributes to these phenotypes, which we have named the group <u>B</u> streptococcal <u>L</u>XG <u>T</u>oxin <u>A</u> (BltA). We demonstrate that BltA is secreted by the T7SS and is toxic to other bacteria when intracellularly co-expressed with associated protein chaperones. Finally, we show that BltA and its associated chaperones contribute to GBS vaginal colonization and ascending infection. Altogether, our findings indicate a role for T7SS toxin-mediated interbacterial competition in GBS mucosal persistence.

## RESULTS

### GBS subtype I T7SSb contributes to interbacterial competition

GBS T7SS promotes persistence within the female genital tract tissues, but it remained unclear whether the GBS T7SS may target other bacteria during niche establishment. The contribution of the T7SSb to interbacterial interactions has been demonstrated in a few Gram-positive organisms, with the breadth and specificity of antagonism differing across genera and species. For example, the *S. intermedius* T7SS has been shown to inhibit a broad range of Gram-positive bacteria (13), while the *E. faecalis* T7SS inhibited only a subset of Gram-positive strains tested (26) and *S. aureus* T7SS inhibition was not observed against *S. epidermidis* (27). To evaluate the ability of GBS to compete with other bacterial species, we grew pure cultures of parental or Δ*essC* mutant CJB111 (predators) and a panel of non-kin bacteria (prey) commonly co-isolated with GBS from human polymicrobial environments, such as the diabetic wound and the vaginal tract. Specified predator and prey bacteria were mixed in a defined ratio and co-cultured statically in liquid media for 24 hours before assessing relative prey viability (13, 26, 28). Similar to previous T7SSb studies performed in other Gram-positive species, we did not observe T7SSb-mediated antagonism against Gram-negative organism, *Escherichia coli* (**Fig. 1A**). The CJB111 T7SS also did not inhibit growth of Gram-positive species, *Staphylococcus aureus*, *Staphylococcus epidermidis*, *Streptococcus pyogenes*, or *Lactobacillus crispatus.* We did, however, observe a significant decrease in *Enterococcus faecalis* (representative strain OG1RF) CFU recovered in the presence of the parental CJB compared to the CJB111Δ*essC* mutant. This GBS competition with *E. faecalis*, but not with other Gram-positive species such as *S. aureus*, was also demonstrated in the CNCTC 10/84 clinical isolate background, which expresses a subtype III T7SS locus (**SFig. 1A**). These competition phenotypes were not due to GBS burdens, as the recovered parental GBS and Δ*essC* mutant CFU were equivalent in any given predator-prey competition pair (**SFig 1B-C, SFig. 2A)**.

**Fig 1.**
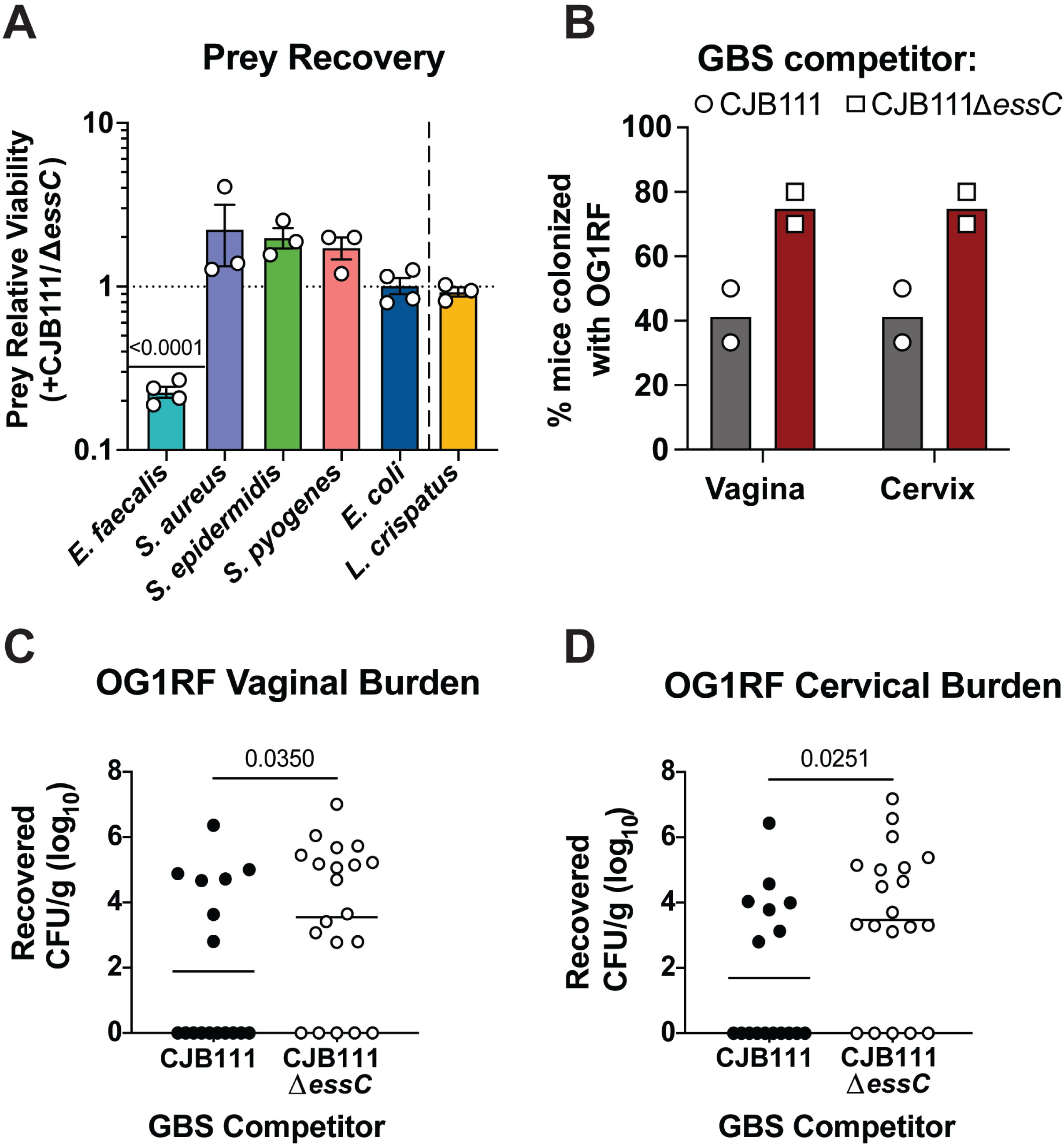
The GBS T7SS reduces recovery of *E. faecalis*, *in vitro* and *in vivo*. **A)** Predator-prey inter-species competition experiments performed in liquid medium between parental CJB111 GBS or CJB111Δ*essC* mutant predator strains and a panel of representative “prey” species. Relative abundance of a given prey strain was calculated as a ratio of prey CFU recovered following 24-hour co-culture with the parental CJB111 predator strain compared to the Δ*essC* predator strain. Statistics reflect one-sample t-tests against a hypothetical value of 1. Data represent the mean of three independent experiments and error bars represent standard error of the mean. **B-D)** GBS T7SS-mediated inhibition of *E. faecalis* OG1RF during murine vaginal colonization. Either parental CJB111 GBS or the CJB111Δ*essC* mutant was co-inoculated with prey *E. faecalis* OG1RF directly into the vaginal lumen (5×10^6^ CFU each strain). **B)** Percent OG1RF colonization of vaginal or cervical tissue plotted for each of two independent experiments (Bars represent the median; Fishers exact test, *p* = 0.0498 in both vaginal and cervical tissue). Recovered *E. faecalis* CFU counts from the **C)** vaginal and **D)** cervical tissues of co-colonized mice. Each dot represents one mouse and data from two independent experiments are combined in these figures (n = 17, 20 total in parental CJB111 and CJB111Δ*essC* mutant co-colonization groups, respectively). The bars in these plots show the median and statistics represent the Mann Whitney U test.

**Fig 2.**
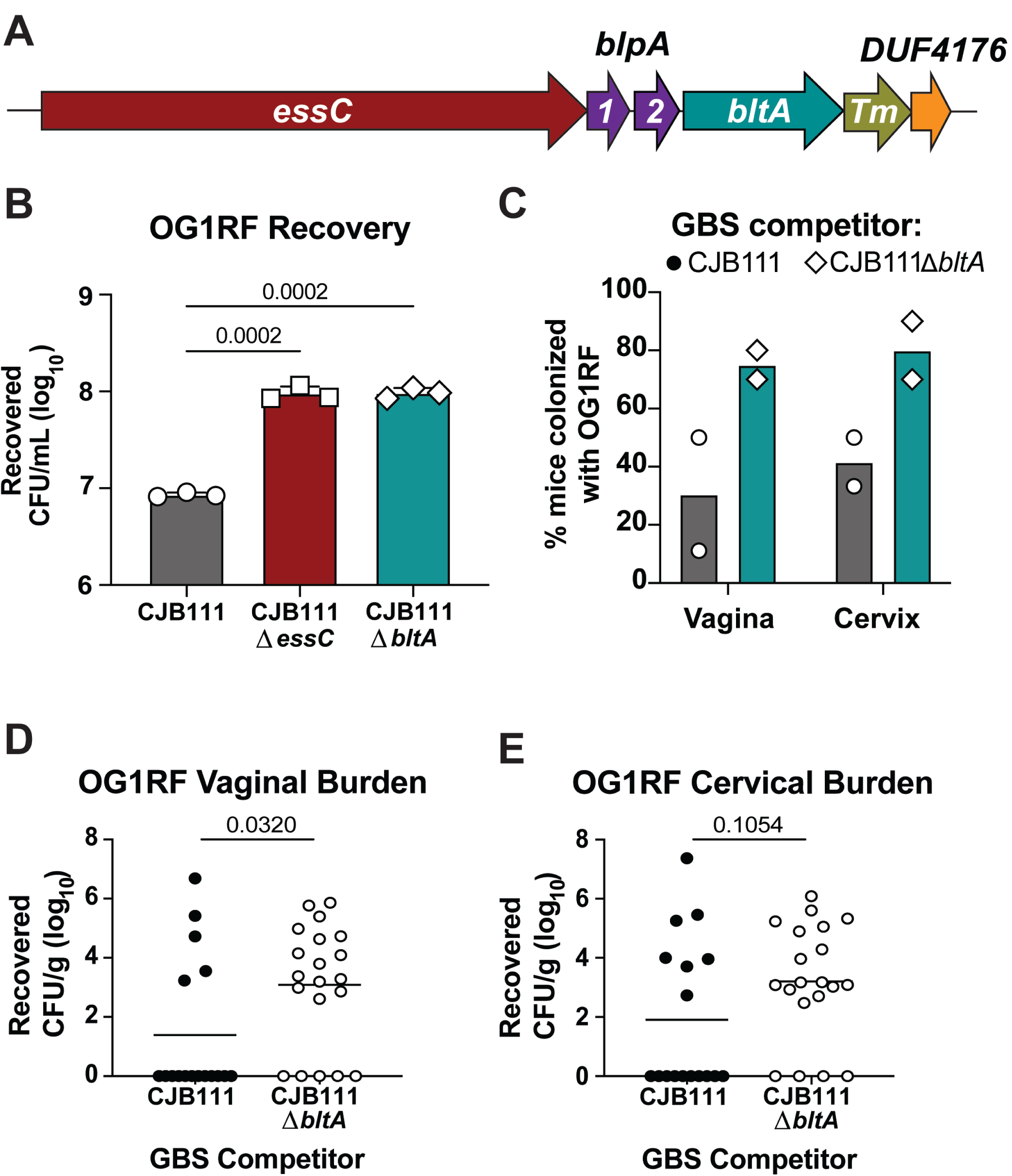
GBS LXG protein BltA contributes to *E. faecalis* antagonism *in vitro* and *in vivo*. **A)** Diagram of part of the GBS T7SS subtype I locus. GBS T7SS loci encode an EssC ATPase ( *essC*), which drives effector export. The subtype I “LXG module” downstream of *essC* includes genes encoding for putative chaperones (BlapA1-2, purple), an LXG toxin (BltA, teal), a Tm protein (olive green), and a DUF4176 protein (orange). **B)** Predator-prey inter-species competition experiments performed in liquid medium between parental CJB111 GBS, CJB111Δ*essC*, or CJB111Δ*bltA* predator strains and *E. faecalis* OG1RF prey. Relative abundance of OG1RF was calculated as a ratio of CFU recovered following 24-hour co-culture with parental CJB111 GBS compared to the CJB111Δ*essC* or CJB111Δ*bltA* mutant strains. Statistics reflect one sample t-tests against a hypothetical value of 1. Data represent the mean of three independent experiments and error bars represent standard deviation. **C-E)** BltA-mediated inhibition of *E. faecalis* during murine vaginal colonization. Either parental CJB111 GBS or the CJB111Δ*bltA* mutant strain was co-inoculated with prey *E. faecalis* OG1RF directly into the vaginal lumen (5×10^6^ CFU each strain). **C)** Percent OG1RF colonization of vaginal or cervical tissue plotted for each of two independent experiments (Bars represent the median; Fishers exact test, *p* = 0.0086 in vaginal tissue and *p* = 0.0210 in cervical tissue). Recovered *E. faecalis* CFU counts from the **D)** vaginal and **E)** cervical tissues of co-colonized mice. Each dot represents one mouse and data from two independent experiments are combined in these figures (n = 17, 20 total in parental CJB111 and CJB111Δ*bltA* mutant co-colonization groups, respectively). The bars in these plots show the median and statistics represent the Mann Whitney U test.

*E. faecalis* is also a vaginal opportunistic pathogen, and our laboratory has previously shown that *E. faecalis* is capable of persisting in murine models of vaginal colonization and ascending infection (29). As the vaginal tract is a common niche for both GBS and *E. faecalis*, we hypothesized that this mucosa may represent a physiologically relevant site of competition between these opportunistic pathogens. To determine if GBS can compete with *E. faecalis* OG1RF in this complex environment, we performed *in vivo* co-colonization experiments in which *E. faecalis* was vaginally co-inoculated with either parental CJB111 or the isogenic CJB111Δ*essC* mutant in conventional mice. We assessed OG1RF tissue burdens at early colonization time points, at which point parental and Δ*essC* mutant GBS burdens remained equal (**SFig. 2B-C**). We found that a higher percentage of mice harbored vaginal and cervical OG1RF when co-colonized with the CJB111Δ*essC* mutant compared to the parental CJB111 (**Fig. 1B**). We further recovered more *E. faecalis* from vaginal and cervical tissue in mice co-colonized with the CJB111Δ*essC* mutant compared to mice co-colonized with parental CJB111 (**Fig. 1C-D)**. Collectively, these data suggest a potential role for the GBS T7SS in competition with another opportunistic pathogen within the physiologically relevant niche of the female genital tract.

### LXG toxin BltA contributes to competition with *E. faecalis*

In other T7SSb+ Gram-positive species, LXG toxins contribute to inhibition of other bacteria. In our previous work, we bioinformatically characterized GBS T7SS loci and found that four subtypes exist that encode for unique putative effector repertoires (24). Downstream of the secretion machinery genes, GBS T7SS subtypes I-III loci encode “modules” containing putative *lxg* genes as well as two putative chaperone-encoding genes (**Fig. 2A**; also known as LXG-associated ɑ-helical proteins; herein called group <u>B</u> streptococcal <u>L</u>XG-associated <u>P</u>roteins, or Blp), which in other species have been shown to facilitate LXG toxin stability and/or secretion (8, 30, 31). Previously, using AlphaFold Multimer modeling, we found that the GBS T7SS subtype I putative chaperones, herein named BlpA1 and BlpA2, are predicted to interact with the putative subtype I LXG toxin, herein named BltA (24). Given the conserved arrangement and function of this module across species, we hypothesized that BltA may promote GBS T7SS interbacterial competition. To assess this, we repeated CJB111 GBS-OG1RF *E. faecalis* predator-prey assays using parental CJB111 and an isogenic CJB111Δ*bltA* mutant. Similar to the Δ*essC* mutant, higher levels of *E. faecalis* were recovered in the presence of the Δ*bltA* mutant compared to the presence of parental CJB111 (**Fig. 2B**), with the GBS CFU recovery remaining equal across these conditions (**SFig. 3A**). To determine if BltA may also contribute to the antagonism against *E. faecalis in vivo,* we repeated the vaginal co-colonization experiments described above, this time co-inoculating either parental CJB111 or the isogenic Δ*bltA* mutant with *E. faecalis* into the vaginal tract. Similar to the results with the Δ*essC* mutant above, we observed that more mice co-colonized with the Δ*bltA* mutant harbored OG1RF within the vagina and cervix (**Fig. 2C**) and at higher tissue burdens (**Fig. 2D-E**) compared to the group co-colonized with parental CJB111. Reciprocally, GBS tissue burdens at this early colonization timepoint did not significantly differ between the groups (**SFig. 3B-C**). These data suggest that BltA contributes to GBS T7SS-mediated interbacterial antagonism against *E. faecalis*.

**Fig 3.**
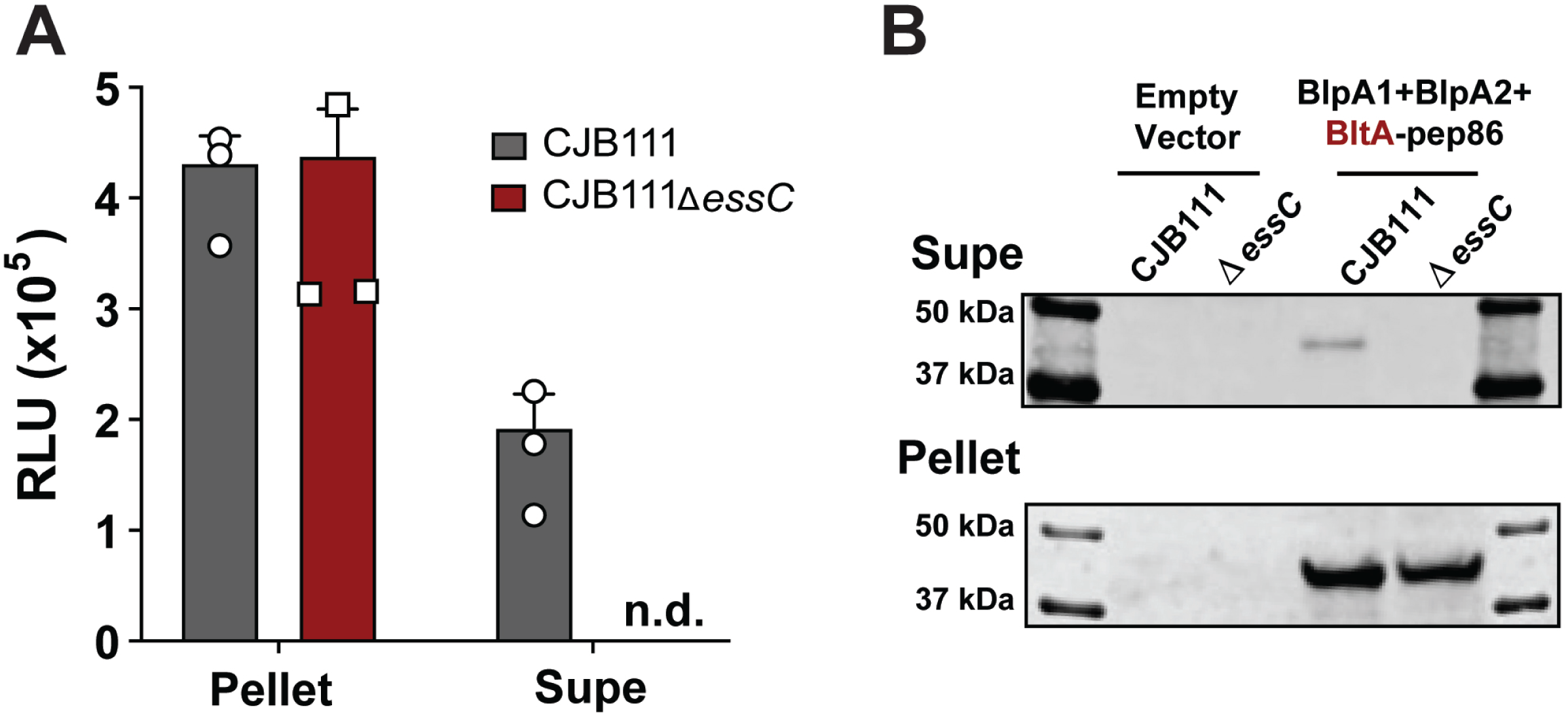
BltA is secreted by the GBS subtype I T7SS in an EssC-dependent manner. **A)** BltA-pep86 was co-expressed with BlpA1-2 chaperones in trans in CJB111 or the CJB111Δ*essC* mutant and luminescence (relative light units, RLU) was detected in cell-associated lysates and cell-free supernatant fractions after addition of the large nanoluciferase 11S fragment and substrate. Data in panel **A** represent the mean of three independent experiments and error bars represent standard deviation. **B)** Western blot showing EssC-dependent secretion of BltA from subtype I strain CJB111. Blot pictured is representative of 3 independent experiments. Coomassie stained gels can be found as loading controls for supernatant and cell-associated pellet fractions in **SFig. 4B**.

### BltA is a secreted substrate of the GBS T7SSb Subtype 1

The *bltA* gene is encoded downstream of the T7SSb machinery genes and adjacent to putative chaperone-encoding genes (*blpA1-2*), a region which typically encodes for T7SSb substrates. To evaluate if BltA is also secreted by the GBS T7SSb, we utilized a Nanoluciferase-based secretion assay (Nanobit) optimized to identify T7SS effectors in *S. aureus* (30, 32, 33). Briefly, the BltA C-terminus was tagged with the small Nanoluciferase subunit pep86 and co-expressed with putative chaperones BlpA1-2 in trans in parental strain CJB111 or in the CJB111Δ*essC* mutant. Secretion of BltA-pep86 was then evaluated by supplementing stationary phase cell lysates or corresponding cell-free supernatants with the large Nanoluciferase 11S subunit (to reconstitute the enzyme) and its substrate and measuring the resulting bioluminescence. Bioluminescent signal exceeding that of the corresponding parental CJB111 or CJB111Δ*essC* mutant empty vector controls (lacking pep86) was reported as relative light units (RLU). While we observed similar luminescence signal from both CJB111 and CJB111Δ*essC* cell-associated lysate fractions, we only observed luminescence from parental CJB111 supernatant, not from CJB111Δ*essC* supernatant (**Fig. 3A**). To confirm BltA secretion by the GBS T7SS, we probed for BltA within these cell lysates and supernatant using an anti-pep86 antibody (Promega). As a control, we also assessed secretion of the canonical T7SS substrate EsxA, which we demonstrated previously to be secreted in an EssC-dependent manner in GBS (**SFig. 4A**) (7). Similar to the Nanobit assay, we observed EssC-dependent BltA secretion by Western blot (**Fig. 3B**) and did not detect any signal in the cell associated or supernatant of the empty vector control strains (which lack the pep86 tag). Collectively, these data indicate that BltA is indeed a GBS T7SSb subtype I substrate.

**Fig. 4.**
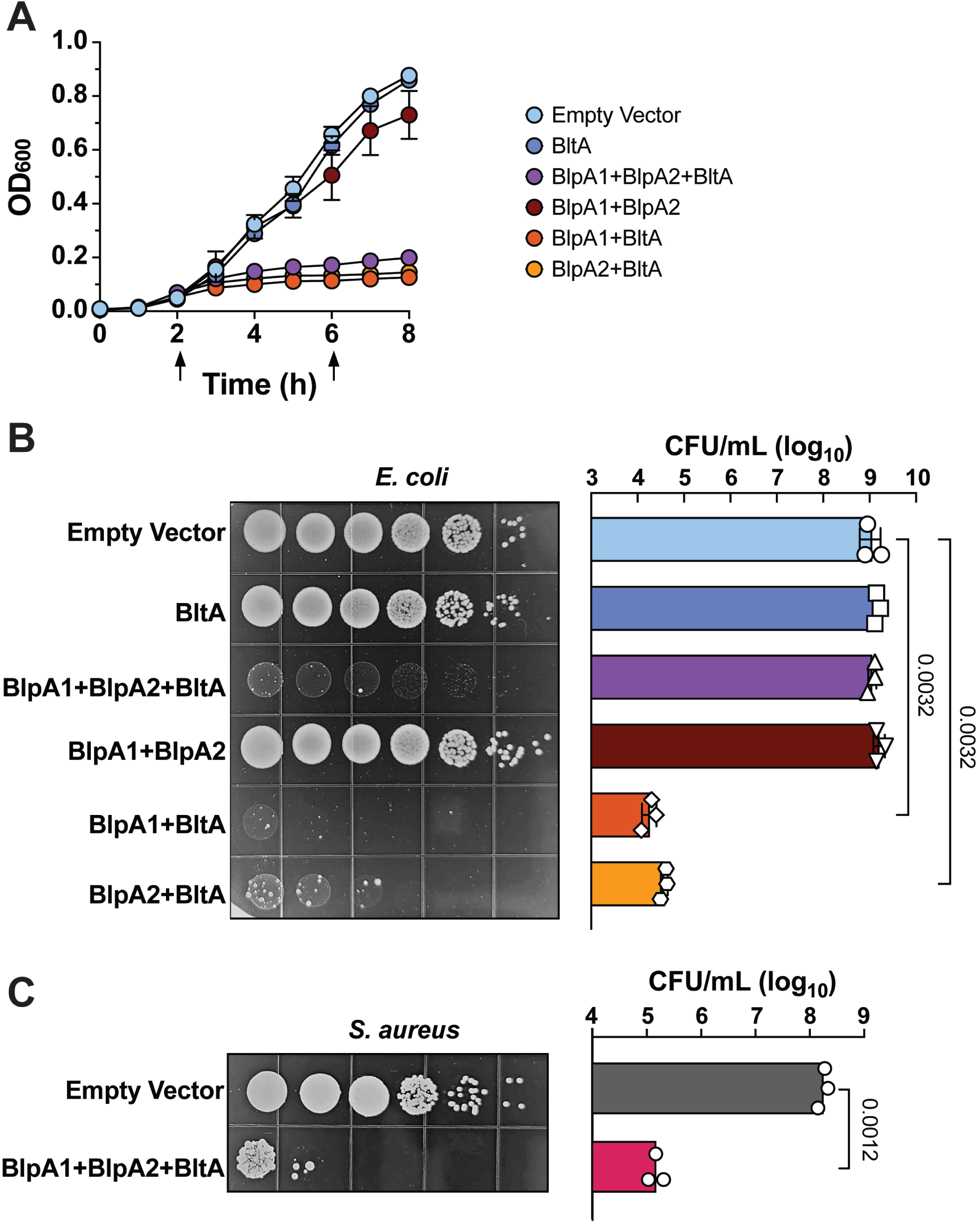
Inducible intracellular expression of BltA with associated chaperones inhibits growth of *E. coli* and *S. aureus.* **A)** Growth of *E. coli* in liquid medium (as measured by OD_600_) expressing rhamnose-inducible plasmids containing an empty vector, BltA alone, BlpA1 + BlpA2 + BltA, BlpA1 + BlpA2 alone, BltA + BlpA1, or BltA + BlpA2. In panel **A**, arrows indicate addition of rhamnose to induce protein expression. Data in panels **A** represent the mean of three independent experiments and error bars represent standard error of the mean. **B)** Viability of the above *E. coli* strains serially diluted and plated on rhamnose-containing media. Serial dilutions are shown in the representative image on the left and quantified in the panel on the right. **C)** Viability of *S. aureus* strains expressing an empty vector or an inducible plasmid containing BlpA1 + BlpA2 + BltA on xylose-containing media. Serial dilutions are shown in the representative image on the left and results from three independent experiments are quantified in the panel on the right. Data in panels **B-C** represent the mean of three independent experiments and error bars represent standard deviation and comparisons in which *p* < 0.05 are displayed. Statistics in panel **B** reflect the ordinary one-way ANOVA with Dunnett correction for multiple comparisons and statistics in panel **C** reflect the student’s t test.

Finally, secretion of T7SS substrates have been shown to be interdependent (34) but it is unknown if export of known GBS T7SS substrate EsxA is impacted by the presence or absence of other T7SS effectors. To determine if loss of BlpA1-2 or BltA impact EsxA secretion, we assessed extracellular EsxA levels in cell-free supernatant. We detected similar levels of EsxA in the cell-associated pellet and supernatant fractions of Δ*blpA1-2 and* Δ*bltA* mutants compared to that of parental CJB111, indicating that neither GBS BltA nor its associated chaperones BlpA1-2 are necessary for EsxA T7SS export (**SFig. 4C)**.

### Intracellular BltA expression intoxicates bacteria in a chaperone-dependent manner

As T7SS-associated LXG proteins have been shown to have toxic activity, we sought to investigate if BltA might be more broadly toxic upon intracellular expression. To evaluate this, we cloned *bltA* into the pSCRhaB2 vector under the control of a rhamnose inducible promoter as performed previously by others (8, 12–14). Induction of the full length BltA or the C-terminal BltA toxin domain alone in liquid culture had no or little impact on *E. coli* growth, respectively (**Fig. 4A, SFig. 5A)**. To assess if chaperones are required for BltA toxicity, we co-expressed *bltA* along with *blpA1*, *blpA2*, or both *blpA1 and blpA2* together. Compared to *E. coli* harboring an empty vector control, we observed a drastic reduction of growth in liquid culture by *E. coli* expressing *blpA1* and *blpA2* with *bltA*, but not upon expression of *blpA1-2* alone **(Fig. 4A)**. Interestingly, we found that co-expression of either *blpA1* or *blpA2* with *bltA* was sufficient for this inhibition of *E. coli* growth **(Fig. 4B)** but that none of these conditions appear to result in lysis (ss measured by OD_600_). To further investigate the impact of BltA on *E. coli* viability, we plated serial dilutions of the above strains on solid media containing or lacking the rhamnose inducer. Similar to the above growth assays, induction of expression of either *bltA* or *blpA1* + *blpA2* alone did not impact *E. coli* CFU recovery. Interestingly, we found that co-expression of either *blpA1* or *blpA2* with *bltA* significantly decreased *E. coli* viability compared to the empty vector control, whereas co-expression of both *blpA1-2* with *bltA* only impacted *E. coli* colony morphology **(Fig 4B)**. As strains plated on solid agar lacking rhamnose (uninduced) exhibited equivalent CFU recovery (**SFig. 5B),** these data suggest that induction of BltA along with a chaperone is toxic to *E. coli*.

**Fig. 5.**
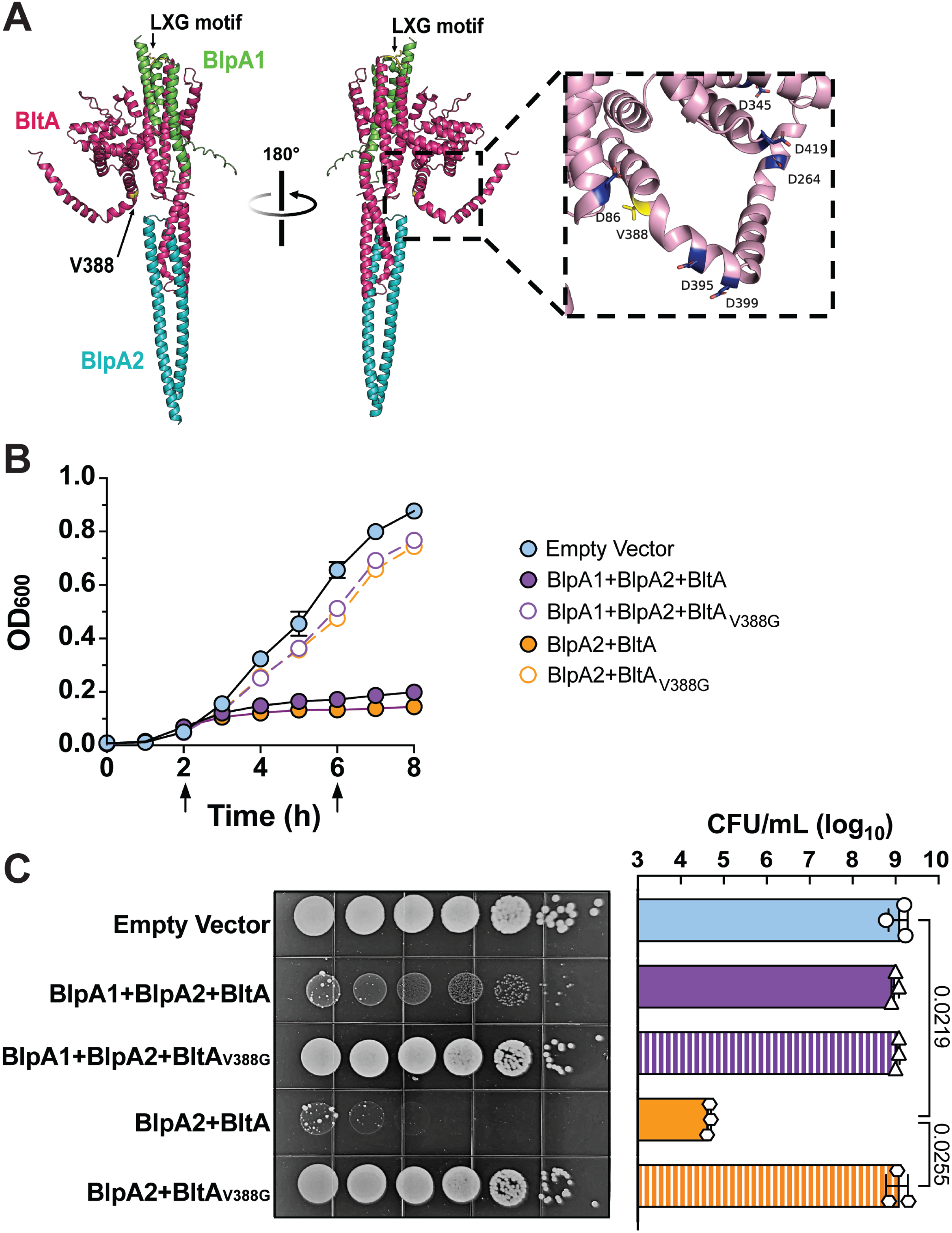
Identification of BltA-resistant *E. coli*. BltA-resistant colonies identified in **Fig. 4** were isolated and their plasmids were sequenced to identify mutations abrogating toxicity. **A**) AlphaFold Multimer predicted model of BltA, BlpA1, and BlpA2 labeled in magenta, green, and blue, respectively. The N-terminal LXG motif and residue V388 are highlighted in yellow. Modelling confidence metrics are presented in **SFig. 6**. Growth of *E. coli* in **B)** liquid or on **C)** solid medium is not impacted by induction of BltA harboring the V388G mutation compared to the empty vector control. CFU recovery of uninduced strains from panel **C** is provided in **SFig 7B**. Data in panels **B-C** represent the mean of three independent experiments and error bars represent standard error of the mean and standard deviation, respectively. Statistics in panel **C** reflect the ordinary one-way ANOVA with Tukey’s multiple comparison test and comparisons in which *p* < 0.05 are displayed.

To investigate whether BltA is also toxic when expressed within Gram-positive bacteria, we next sought to assess BltA inhibition of *S. aureus,* which was also not susceptible to GBS T7SS in direct competition experiments (**Fig. 1A)**. To evaluate this, we cloned *blpA1-2* and *bltA* into the pEPSA5 expression vector under the control of a xylose-inducible promoter and plated serial dilutions on solid media containing or lacking the xylose inducer. We found that induction of *blpA1-2 + bltA* intracellular expression resulted in approximately 1000-fold decreased CFU recovery of *S. aureus* compared to the empty vector **(Fig. 4C)** and uninduced controls (**SFig. 5C**). These data suggest that BltA is capable of intoxicating both Gram-negative and Gram-positive hosts intracellularly and that its target is likely conserved across genera.

### BltA toxicity is abrogated by a missense mutation within the C-terminal putative toxin domain

In performing the above inducible intoxication assays, *bltA* expression along with either *blpA1* or *blpA2* resulted in a dramatic reduction in *E. coli* viability. Interestingly, in these assays, we often observed resistant mutants that were unaffected by *bltA* induction **(Fig. 4B)**. We isolated single resistant colonies and sequenced their plasmids to identify potential mutations that may impact BltA’s toxic activity. We found many mutations across these plasmids, including large insertions within *bltA* or its associated chaperones, a slip-strand mutation within *bltA* resulting in a frameshift, as well as a missense V388G mutation within the putative *bltA* toxin domain (**SFig. 7A**, **Fig 5A**). Induction of *bltA*_V388G_ co-expressed with *blpA2* within a clean *E. coli* background (to exclude any additional chromosomal mutations) significantly reduced BltA inhibition in both liquid and agar media (**Fig. 5B-C**). Further, *bltA*_V388G_ co-expressed with both *blpA1-2* reduced BltA growth inhibition in liquid agar and restored the altered *E. coli* morphology observed in cells expressing *blpA1-2* + *bltA*. These data suggest that this V388 residue may impact BltA function.

### BltA and BlpA1-2 chaperones promote GBS vaginal colonization

Previously we showed that the T7SS is important for colonization of the vaginal tract by subtype I-expressing strain CJB111. However, the T7SS effectors that mediate this colonization were not investigated. Here we sought to determine if BltA might impact GBS colonization. To investigate this, we performed single challenge vaginal colonization experiments inoculating either parental CJB111 or the isogenic CJB111Δ*bltA* mutant into the vaginal tract and assessed GBS persistence in female genital tract tissues. Similar to those observed with our T7SS-deficient CJB111Δ*essC* mutant (24), we observed differences in tissue burdens in the vagina and cervix between mice colonized with the CJB111Δ*bltA* mutant compared to parental CJB111 (**Fig. 6A-B**). As the BlpA1-2 chaperones appear to be important for BltA toxicity, we further hypothesized that loss of these chaperones might also impact GBS persistence within female genital tract tissues. Indeed, upon GBS single-challenge into the vaginal tract, we observed that the CJB111Δ*blpA1-2* mutant was recovered in significantly lower burdens from the vagina and cervix compared to parental CJB111 (**Fig. 6C-D**). These results further demonstrate the importance of BltA and its associated chaperones during *in vivo* GBS colonization.

**Fig. 6.**
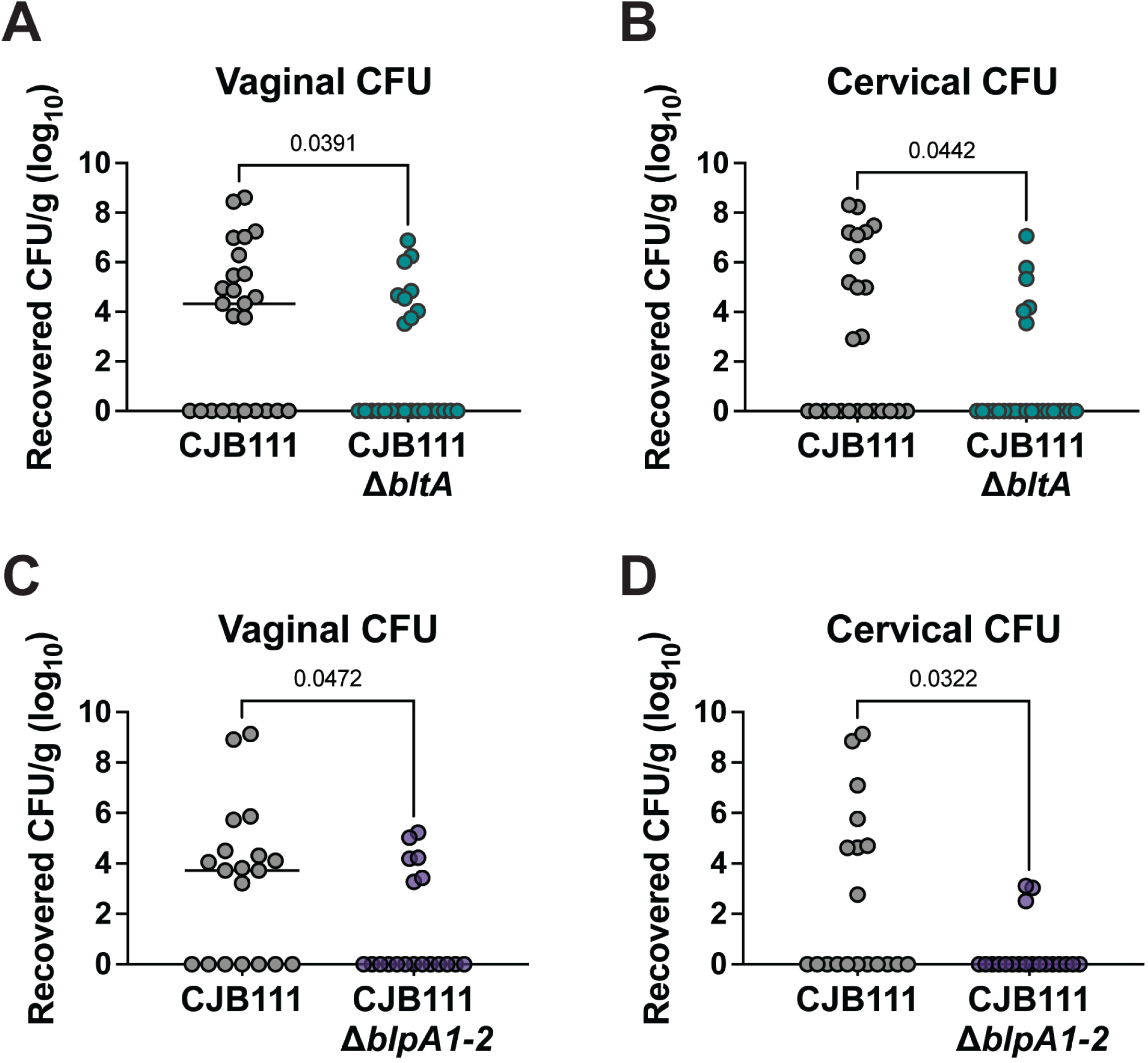
Loss of BltA or associated chaperones impacts GBS vaginal colonization by T7SS subtype I strain CJB111. Recovered CFU counts from the **A, C)** vaginal and **B, D)** cervical tissue of mice mono-colonized with parental CJB111 GBS and either the CJB111Δ*bltA* or CJB111Δ*blpA1-2* mutant strains. Each dot represents one mouse and all independent experiments’ data are combined in these figures (n = 25/group for Δ*bltA* experiments and n = 19/group for Δ*blpA1-2* experiments). The bars in these plots show the median and statistics represent the Mann Whitney U test.

## Discussion

Herein, we demonstrate that GBS T7SS reduces recovery of mucosal pathobiont *E. faecalis*, both *in vitro* and *in vivo* within the complex environment of the murine female genital tract. We have further identified BltA as an LXG toxin encoded downstream of T7SS machinery in GBS subtype I that contributes to this competition and which is toxic to other bacteria upon intracellular expression. Investigation of resistant *E. coli* revealed mutations within the chaperones and BltA, including a BltA V388G missense mutation that abrogates toxicity. Finally, we show that BltA and its associated chaperones promote GBS persistence within the vagina and cervix during colonization.

In this work, we primarily examined GBS in competition with other pathobionts such as *Enterococcus faecalis*, another vaginal opportunistic pathogen (29) that may concurrently colonize the female genital tract during conditions such as aerobic vaginitis (35, 36). We observed T7SS-mediated *E. faecalis* antagonism by two different GBS clinical isolates. Interestingly, under the conditions used here, we did not observe GBS T7SS subtype I-mediated competition with human vaginal commensal *Lactobacillus crispatus* and instead observed a T7SS-independent decrease in GBS CFU recovery following co-culture. Consistently, lactobacilli have been shown previously to impact GBS virulence and viability and have therefore been suggested as a potential therapeutic to reduce or prevent GBS colonization of the vaginal mucosa (21, 37, 38). While future work will investigate the interaction of the GBS T7SS with other vaginal commensals, it is possible that the T7SS may have evolved to target other pathobionts to promote niche establishment rather than members of the native vaginal microbiota. However, this remains to be determined. Importantly, while T7SSb-mediated interbacterial antagonism has been demonstrated for several Gram-positive T7SSb+ organisms, the conditions used during predator-prey assays likely influence whether the T7SSb is induced and therefore whether LXG toxin-mediated competition is observed or not (27, 39). Therefore, identifying factors and conditions that induce GBS T7SSb expression is part of our ongoing work.

We further show here that induction of intracellular BltA expression is toxic to both *S. aureus* and *E. coli.* While no obvious conserved domain or motifs exist within the BltA C-terminal toxin domain to inform its function, based on the above data, the target of this toxin appears to be intracellular and conserved across Gram-positive and Gram-negative bacteria. Because neither *S. aureus* nor *E. coli* was inhibited by GBS during predator prey assays, it is possible that an extracellular factor may be required for T7SS-mediated antagonism. To date, it is unclear how T7SSb effectors, including LXG toxins, are delivered into a prey cell, although a few studies have indicated that T7SSb competition is contact-dependent (11, 13, 26). We hypothesize that specific extracellular determinants may dictate the range of GBS T7SS antagonism and will investigate this in our future work.

During inducible intoxication assays in *E. coli*, we found that BltA alone was not sufficient to reduce bacterial viability but that co-expression of BlpA1 and/or BlpA2 promoted BltA intoxication in liquid medium. Further, some of the mutations we identified in resistant colonies disrupted the BlpA putative chaperones or the N-terminal LXG domain of BltA that is predicted to interact with these chaperones. This is consistent with previous literature that Laps may promote LXG stability. Interestingly, co-expression of a single chaperone with BltA resulted in decreased *E. coli* viability during solid agar growth assays compared to expression of BltA with both BlpA1 and BlpA2, indicating that specific chaperone interactions may impact BltA toxicity. Future work will investigate the molecular interactions of BlpA1 and Blp2A with BltA as well as the role of each putative chaperone in secretion, stability, and/or toxin function.

In addition to mutations in BlpA1 and BlpA2, a V388G mutation was identified in two independent experiments within the BltA putative toxin domain. As co-expression of BltA_V388G_ with its chaperone(s) is not toxic, it is possible that the V388G mutation may disrupt the stability and/or function of BltA. Although this region of the model is predicted by AlphaFold at very low confidence, this V388G mutation occurs within the C-terminal putative toxin domain of BltA and is proximal to a several aspartate residues in multiple predictive iterations (**Fig. 5A**). Aspartate rich regions are associated with some enzyme active sites and an aspartate-rich motif has been identified previously within the *Streptococcus intermedius* LXG toxin TelC to be important for its function (13). Whether BltA functions as an enzyme is still unknown and our future work will investigate whether these aspartate residues are important for BltA toxicity and how mutations in this area impact the structure and function of BltA.

While in this study we have focused on T7SSb effectors encoded by the highly prevalent GBS T7SS subtype I, other subtypes of GBS T7SS remain unstudied. GBS T7SS subtypes I - III encode similar types of putative effectors and accessory proteins, including LXG toxins, associated chaperones, transmembrane (Tm) proteins, and DUF4176-containing proteins. Despite this conservation, these protein products are largely unique between subtypes at the sequence level. Indeed, while each putative LXG toxin from subtypes I-III contains the conserved N-terminal LXG domain, each encodes a unique C-terminal putative toxin domain. We have observed previously that none of these LXG toxins contain easily identifiable functional domains; thus, the biochemical activity of these proteins must be determined experimentally. We observed that the subtype I LXG toxin BltA contributes to *E. faecalis* antagonism and, given we also observed T7SS subtype III-dependent *E. faecalis* inhibition by clinical isolate CNCTC 10/84, future work will investigate potential contribution of the subtype III LXG protein to competition as well as the potential antagonistic range of the additional GBS T7SS subtypes II and IV.

Toxins across many specialized secretion systems commonly encode an immunity factor nearby to prevent self-intoxication. Indeed, in other bacterial species in which LXG toxins have been studied, a cognate immunity factor was identified either one or two genes downstream of the toxin encoding gene. Within the GBS subtype I locus, *bltA* followed by two genes of unknown function. We first scanned other bacterial genomes for BltA homologs to evaluate whether a common gene is always encoded downstream, which would strongly indicate its function as an immunity factor. Upon BLAST search of BltA in non-GBS genomes, we identified a homolog (81% identical) in mastitis pathogen *Streptococcus uberis* (GenBank accession SAMN27553288). This *S. uberis* locus encodes an LXG module syntenic to that found in the subtype I GBS, including homologs of BlpA1 (72% identical) and BlpA2 (41% identical) encoded upstream of BltA. The *S. uberis* locus downstream of *bltA* also encodes for a Tm-encoding gene as well as a DUF4176 domain containing protein (85% identical), similar to those found in GBS subtype I. Interestingly, the *S. uberis* Tm-encoding gene contains two slip-strand mutations resulting in frameshift and premature truncation of the protein (resulting in correct translation of 115 of 182 amino acids). Because of this, we hypothesize that this Tm protein may not be the immunity factor, although it is possible that the truncated Tm protein is sufficient to prevent intoxication. Alternatively, BltA may be tightly regulated as to not require an endogenous downstream immunity factor or another downstream gene may encode for the BltA immunity factor. This comprises a direction of our future work.

To conclude, in this work, we have demonstrated GBS T7SS-dependent inhibition of *E. faecalis in vitro* for two distinct T7SS subtypes as well as T7SS subtype I-dependent *E. faecalis* antagonism in the murine female genital tract *in vivo*. We identified a novel GBS T7SS LXG effector, BltA, as a mediator of this interbacterial antagonism. Our work further indicates a role for BltA as an antibacterial toxin with an intracellular target conserved across Gram-positive and Gram-negative genera. As has been seen in multiple streptococcal species as well as *S. aureus*, we observed that two putative ɑ-helical chaperone proteins, BlpA1 and BlpA2, are important for BltA toxicity. Future work will characterize the mechanisms driving BltA toxicity including the function of BlpA1 and BlpA2, the target of BltA, and identifying the cognate immunity factor. Additionally, we seek to identify prey bacterial factors that contribute to the breadth and specificity of GBS T7SS-mediated interbacterial antagonism against not only vaginal pathobionts, but those found in broader GBS host polymicrobial niches such as the neonatal gastrointestinal tract as well as in diabetic wounds.

## Materials/Methods

### Bacterial strains

Bacterial strains used in this study can be found in **Supplemental Table 1**. GBS strains were grown statically in Todd Hewitt broth (THB; Research Products International, RPI Catalog# T47500) at 37°C with erythromycin (5 μg/mL) when applicable. *E. coli* strains were grown shaking in Luria broth (LB; RPI Catalog# L24066) at 37°C supplemented with erythromycin (500 μg/mL), carbenicillin (100 μg/mL), trimethoprim (200 μg/mL), or L-rhamnose (0.1% w/v) when applicable. *S. aureus* strains were grown shaking in Tryptic soy broth (TSB; Becton Dickinson, BD Catalog# 211822) at 37°C supplemented with chloramphenicol (10 μg/mL) or xylose (2% w/v) when applicable. *E. faecalis* was grown shaking in THB at 37°C. *S. epidermidis* was grown shaking in TSB at 37°C. *S. pyogenes* was grown statically in THB at 37°C in 5% CO_2_. *L. crispatus* was grown statically in de Man, Rogosa, Sharpe broth (MRS; ThermoFisher Scientific Catalog #CM0361) at 37°C in 5% CO_2_.

### Cloning

Clean *bltA and blpA1-2* deletion mutants were created via allelic exchange in CJB111 using the temperature sensitive plasmid pHY304 and using a gene encoding spectinomycin resistance ( *aad9*) in the knockout construct. Second crossover mutants were screened for erythromycin sensitivity and spectinomycin resistance. For intoxication assays, genes encoding for *blpA1, blpA2,* and/or *bltA* were amplified from the CJB111 and ligated into the inducible *E. coli* expression vector pSCRhaB2 or the inducible *S. aureus* expression vector pEPSA5 via Gibson assembly. Constructs for the Nanobit nanoluciferase assay were generated by adding a pep86 tag and short linker sequence (provided by Promega) to the 3’ end of *bltA* using Phusion polymerase. Amplicons were then ligated into the GBS expression vector pDCErm using Quick Ligase (restriction enzymes BamHI-HF and SacII-HF) or Gibson assembly. Strains created and primers used in this study can be found in **Supplemental Table 1.**

### Interbacterial Competition

GBS predator and designated prey strains were grown overnight for 16-20 hours under the standard laboratory conditions listed above for each strain. Except for *L. crispatus,* all prey co-cultures were prepared as follows: for bacteria that were not grown overnight in THB, cells were first pelleted, the supernatant discarded, and finally resuspended in fresh THB. All strains were then normalized to OD_600_ = 0.4 in THB. Prey strains were diluted 1:1000 in THB. 10-fold serial dilutions of each prepared predator and prey culture were plated on respective standard agar media and incubated under respective standard laboratory conditions to confirm predator-prey CFU ratios. Competition cultures were prepared by mixing 1 mL of normalized GBS predator culture with 1 mL of prepared prey culture in a 5 mL culture tube (Falcon, Catalog# 352054). Cultures were then vortexed briefly before being incubated statically at 37°C, 5% CO_2_ for 24 hours. Monoculture controls were included wherein each prepared culture was mixed with 1 mL THB rather than a second strain. After 24 hours, serial dilutions of each co-culture were plated on differential and/or selective media to assess final CFU of both GBS and the designated prey strain. Medias used were as follows: CHROMagar™ StrepB (CHROMagar Catalog# SB282) for GBS; THA containing rifampin (50 μg/mL) and fusidic acid (25 μg/mL) for *E. faecalis;* mannitol salt agar (BD, Catalog# 211407) for *S. aureus* and *S. epidermidis*; THA and THA containing tetracycline (5 μg/mL) for *S. pyogenes* (CFU recovered from THA with tetracycline were subtracted from CFU recovered from THA to calculate *S. pyogenes* CFU); and MacConkey agar (BD Catalog# 212123) for *E. coli*. Monocultures were plated on the same media as the input cultures. All plates were then incubated 16-24 hours at 37°C (with 5% CO_2_ for *S. pyogenes*). CFU was enumerated for each strain and prey relative viability was calculated as a ratio of the prey CFU recovered from co-culture with parental GBS compared to co-culture with mutant GBS. This competition protocol was slightly modified for *L. crispatus* competition due to inherent lactobacilli growth constraints. Rather than normalizing and incubating strains in THB, both GBS and *L. crispatus* were prepared and incubated in MRS. Control experiments were performed to confirm there was no growth defect of GBS in MRS (data not shown). Monocultures and co-cultures were set up as described above. Both *L. crispatus* monocultures and co-cultures were plated on MRS agar and incubated in 5% CO_2_. *L. crispatus* CFU was enumerated, and relative viability was then calculated as above.

### Vaginal colonization

GBS vaginal colonization was assessed using a previously described murine model of vaginal persistence and ascending infection (40). Briefly, 8–10-week-old female CD1 (Charles River) mice were synced with beta-estradiol at day −1 and inoculated intravaginally with mid-log grown GBS (approximately 1 × 10^7^ CFU) in PBS on day 0. Post-inoculation, mice were lavaged with PBS daily, and the samples were serially diluted and plated for CFU counts to determine bacterial persistence on CHROMagar. At experimental end points, mice were euthanized, and female genital tract tissues (vagina and cervix) were collected. Tissues were homogenized and samples were serially diluted and plated on CHROMagar for CFU enumeration. Bacterial counts were normalized to the tissue weight. Co-challenge GBS-*E. faecalis* vaginal colonization was performed as described above, except that mice were co-challenged with approximately 5 × 10^6^ CFU *E. faecalis* + 5 × 10^6^ CFU GBS. Lavage and tissues were serially diluted and plated on CHROMagar for GBS enumeration and THA containing rifampin (50 μg/mL) and fusidic acid (25 μg/mL) for *E. faecalis* enumeration. Vaginal colonization experiments were ended upon vaginal clearance of one strain by 70% of the mice. Mice not initially colonized or those recovering maximal GBS colonization following multiple days of luminal clearance were excluded. These experiments were approved by the committee on the use and care of animals at the University of Colorado-Anschutz Medical Campus (protocol #00316).

### Detection of cell-associated and secreted T7SS effectors by western blot

Secretion of EsxA and BltA during laboratory growth was detected as described previously (7), with slight modifications for strains expressing pep86 tagged effectors. Briefly, GBS strains harboring pep86-tagged constructs and respective empty vector strains were grown with the addition of erythromycin (5 μg/mL) to the culture media. Stationary phase GBS cultures were separated into pellet and supernatant fractions via centrifugation. Supernatants were filtered (Millex Low Protein Binding Durapore PVDF Membrane 0.22μm filters, catalog #SLGVR33RS), supplemented with an EDTA-free protease inhibitor cocktail (Millipore-Sigma set III, catalog # 539134; 1:250 dilution), and proteins were precipitated with trichloroacetic acid (resuspended in Tris buffer (50 mM Tris HCl, 10% glycerol, 500 mM NaCl, pH 7)). Bacterial pellets were lysed by bead-beating in Tris buffer + protease inhibitor and supplemented with Triton-X-100 to solubilize membrane proteins. Supernatant and pellet fractions were run on SDS-PAGE and stained with Coomassie stain or transferred to membranes as described previously. Membranes were blocked in LI-COR blocking buffer and then probed with anti-HiBiT (pep86) mouse monoclonal antibody (1 μg/ml; Promega, Catalog# N7200) and/or anti-EsxA1 rabbit polyclonal antibody (0.5 μg/ml; GenScript) in LI-COR blocking buffer overnight at 4°C. Following washes, membranes were incubated with IRDye 800CW goat anti-mouse IgG and/or IRDye 680RD goat anti-rabbit IgG secondary antibodies from LI-COR (1:10,000 dilution; 1 hour, room temperature). Following washes in TBST and water, western blots were imaged using the LI-COR Odyssey.

### Detection of cell-associated and secreted T7SS effectors by Nanobit assay

GBS strains harboring pep86-tagged constructs and respective empty vector strains were grown as in the western blot protocol above, with the addition of erythromycin (5 μg/mL) to the culture media. Pellet and supernatant fractions were prepared as above. Pellet and concentrated supernatant samples were diluted 1:2 or 1:10, respectively, in Tris buffer. Then, 100 μL of each sample were added to a 96-well white microplate (Costar). The pep86-tag was detected in each fraction using the Nano-Glo® HiBiT Extracellular Detection System (Promega Catalog# N2420) according to the manufacturer’s instructions. Luminescence was measured with a Tecan Infinite M Plex plate reader. Relative luminescence from BltA-pep86 expressing parental CJB111 or CJB111Δ*essC* strains were normalized to their respective empty vector controls.

### Bacterial Intoxication Assays

*E. coli* strains containing pSCRhaB2 constructs were grown shaking overnight in LB with trimethoprim (200 μg/mL) at 37°C. Each strain was then normalized to OD600 = 1 before being diluted 1:100 into fresh LB with trimethoprim (200 μg/mL). For OD600 growth curves, 150 μL of each strain were added to a 96-well microtiter plate in technical triplicate. Plates were grown shaking at 800 rpm, 37°C and OD600 was measured every hour with a Tecan Infinite M Plex plate reader. Genes expressed on pSCRhaB2 were induced at 2 and 6 hours with 0.1% w/v L-rhamnose (or at 2, 4, 6, and 8 hours for the C-terminal BltA construct). Uninduced controls were included for each experiment. For agar toxicity assays, OD600 normalized cultures from above were serially diluted 10-fold to 10^-8^ in PBS and 10 μL of each dilution was spotted onto LB agar plates containing trimethoprim (200 μg/mL) with or without 0.1% w/v L-rhamnose. Plates were then incubated 16 hours at 37°C before imaging and enumerating CFU. *S. aureus* pEPSA5 strains were grown shaking overnight in TSB with chloramphenicol (10 μg/mL) at 37°C. Each strain was then normalized to OD600 = 0.5 before being serially diluted 10-fold to 10^-8^ in PBS. 10 μL of each dilution was spotted onto TSA containing chloramphenicol (10 μg/mL) with or without 2% w/v xylose. Plates were incubated 16 hours at 37°C before imaging and enumerating CFU.

### AlphaFold modeling

Predictive modelling of the BlpA1-BlpA2-BltA complex was performed using AlphaFold2-multimer weights (41) and implemented in ColabFold (42). Illustrations were generated using PyMOL (The PyMOL Molecular Graphics System, Version 2.5.5, Schrödinger, LLC) (43).

### Statistics

Statistical analysis was performed using Prism version 10.2.3 for macOS (GraphPad Software, La Jolla, CA, United States). Significance was defined as p < α, with α = 0.05.

## Supplemental Figure Legends

**SFig. 1. GBS isolate CNCTC10/84 displays a similar interbacterial antagonism profile to CJB111**

**A)** Predator-prey interbacterial competition assay performed between parental or Δ*essC* mutants of GBS CJB111 or CNCTC 10/84 and prey *E. faecalis* or *S. aureus.* Prey relative viability is calculated as in **Fig. 1A**. Statistics reflect one-sample t-tests against a hypothetical value of 1. Data represent the mean of three independent experiments and error bars represent standard deviation. Parental or Δ*essC* mutant GBS CFU recovery for **B)** CJB111 and **C)** CNCTC 10/84 strains following *in vitro* predator-prey inter-species competition experiments performed in **SFig. 1A**. Statistics reflect multiple unpaired t-tests. Data represent the mean of three independent experiments and error bars represent standard deviation.

**SFig. 2. Parental CJB111 and CJB111Δ*essC* CFU are recovered equally upon co-culture with a given prey during *in vitro* and *in vivo* competition assays. A)** Parental CJB111 and CJB111Δ*essC* mutant CFU recovery following *in vitro* predator-prey inter-species competition experiments performed in **Fig. 1A**. Statistics reflect multiple unpaired t-tests. Data represent the mean of three independent experiments and error bars represent standard deviation. GBS CFU burden recovered from the **B)** vagina and **C)** cervix during *in vivo* co-colonization competition experiments performed in **Fig. 1C-D**. Each dot represents one mouse and data from two independent experiments are combined in these figures (n = 17, 20 total in parental and Δ*essC* mutant co-colonization groups, respectively). The bars in these plots show the median and statistics represent the Mann Whitney U test.

**SFig. 3. GBS CFU recovery during *in vitro* and *in vivo* competition assays using the Δ*bltA mutant*.**

**A)** Parental CJB111, CJB111Δ*essC* mutant, and CJB111Δ*bltA* mutant CFU recovery following *in vitro* predator-prey inter-species competition experiments performed in **Fig. 2B**. Statistics reflect multiple unpaired t-tests. Data represent the mean of three independent experiments and error bars represent standard deviation. GBS CFU burden recovered from the **B)** vagina and **C)** cervix during *in vivo* co-colonization competition experiments performed in **Fig. 2D-E**. Each dot represents one mouse and data from two independent experiments are combined in these figures (n = 17, 20 total in parental and Δ*bltA* mutant co-colonization groups, respectively). The bars in these plots show the median and statistics represent the Mann Whitney U test.

**SFig. 4. Impact of EsxA secretion upon overexpression of BltA-pep86 or loss of BltA or BlpA1-2.**

**A)** Western blot showing EssC-dependent secretion of EsxA from subtype I CJB111 strains overexpressing BlpA1+BlpA2+BltA-pep86 compared to CJB111 empty vector controls. **B)** Coomassie-stained SDS PAGE gel indicating that wells were equally loaded for the western blots shown in **Fig. 3B** and **SFig. 4A. C)** Western blot showing equivalent EsxA secretion from CJB111, CJB111Δ*bltA*, and CJB111Δ*blpA1-2* strains. Blots pictured are representative of 3 independent experiments.

**SFig. 5. Controls for BltA intoxication experiments in *E. coli* and *S. aureus* A)** Growth in liquid medium (as measured by OD_600_) of *E. coli* expressing an empty vector control or an inducible plasmid containing the putative toxin domain of BltA. Arrows indicate addition of rhamnose inducer for the induced group. Data represent the mean of three independent experiments and error bars represent standard error of the mean. Growth of **B)** *E. coli* and **C)** *S. aureus* intoxication strains from **Fig. 4C-D** on solid agar media not containing inducer. Serial dilutions are shown in the representative image on the left and results from three independent experiments are quantified in the panel on the right. Data represent the mean of three independent experiments and error bars represent standard deviation. Statistics in panel **C** reflect the student’s t test.

**SFig. 6. Confidence scores and predicted aligned error for BlpA1-BlpA2-BltA AlphaFold predicted models in Fig. 5A. A)** Predicted aligned error for the BlpA1 + BlpA2 + BltA complex. Colors indicate the confidence of domain positions (higher predicted error in red, lower predicted error in blue). Chain numbers A, B, and C correspond with BlpA1, BlpA2, and BltA, respectively. **B)** Predicted BlpA1-BlpA2-BltA complex model with colors corresponding to per-residue confidence level (predicted local distance difference test [pLDDT] score 1-100; pLDDT > 90 are expected to be modelled to high accuracy; pLDDT < 50 may indicate unstructured/disordered regions but should not be interpreted with confidence).

**SFig. 7. Mutations identified in BltA-resistant *E. coli* colonies. A)** Diagram of representative mutations identified in BltA-resistant *E. coli* colonies from strains carrying plasmids encoding BlpA1 + BlpA2 + BltA, BlpA1 + BltA, or BlpA2 + BltA. Mutations included insertion of IS1 and IS3 elements within all three genes on both the forward and reverse strands, a single nucleotide deletion in *bltA*, resulting in premature truncation of the BltA protein, a slip-strand insertion in *bltA* within in a poly-A tract resulting in frameshift, and a BltA V388G missense mutation. Columns indicate the background in which resistant mutations were found. **B)** Growth of *E. coli* intoxication strains from **Fig. 5C** on solid agar media not containing inducer. Serial dilutions are shown in the representative image on the left and results from three independent experiments are quantified in the panel on the right. Data represent the mean of three independent experiments and error bars represent standard deviation.

## Data availability

All relevant data are within the manuscript and its Supporting Information files.

## Ethics statement

Animal experiments were approved by the Institutional Animal Care and Use Committee at University of Colorado-Anschutz protocol #00316 and were performed using accepted veterinary standards. The University of Colorado-Anschutz is AAALAC accredited; and its facilities meet and adhere to the standards in the “Guide for the Care and Use of Laboratory Animals”.

