## Supplementary figures and images for "A group B streptococcal type VII secreted LXG toxin mediates interbacterial competition and colonization of the female genital tract"

### Supplemental Figures

Supplemental Figure 1

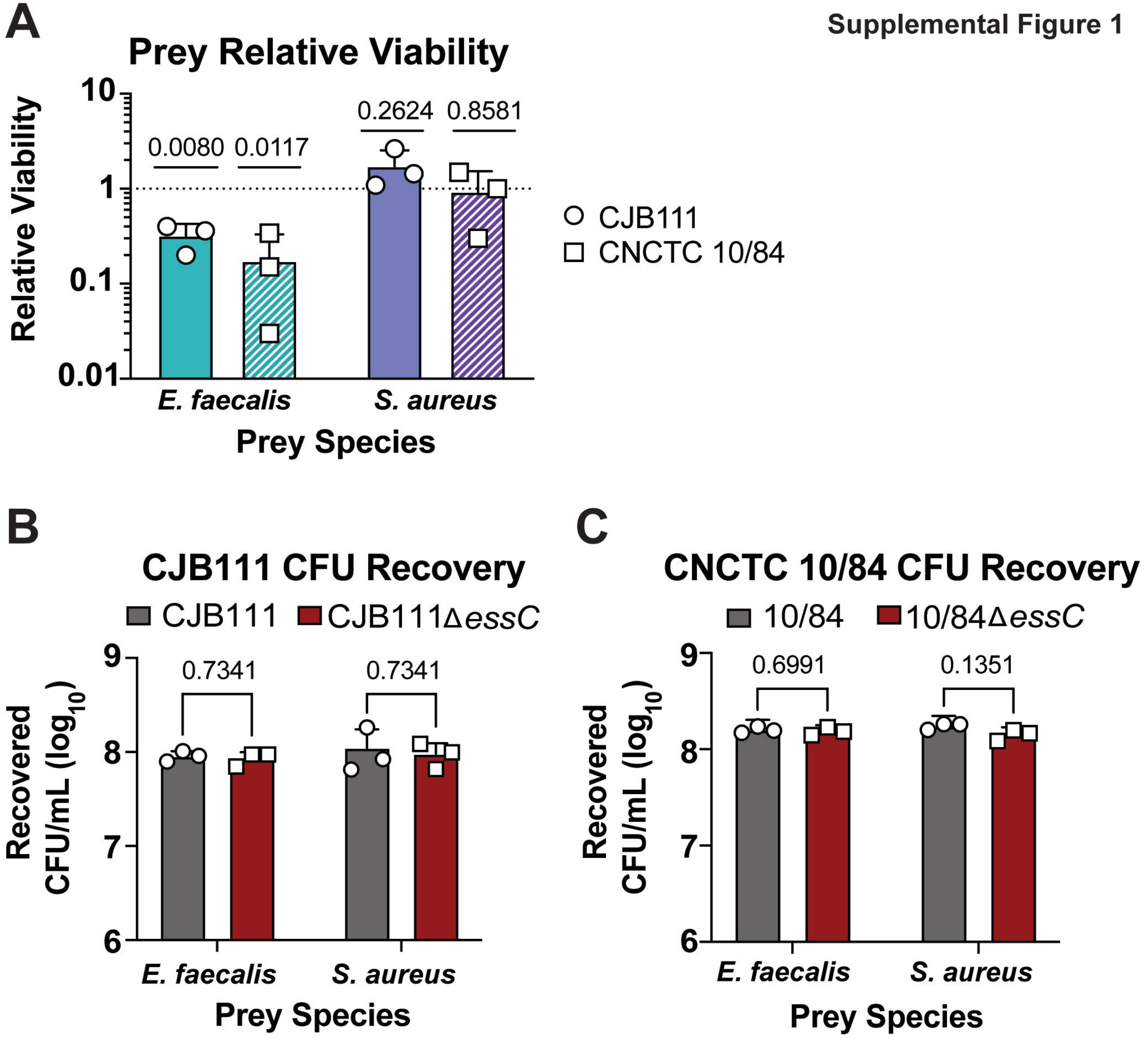

Supplemental Figure 2

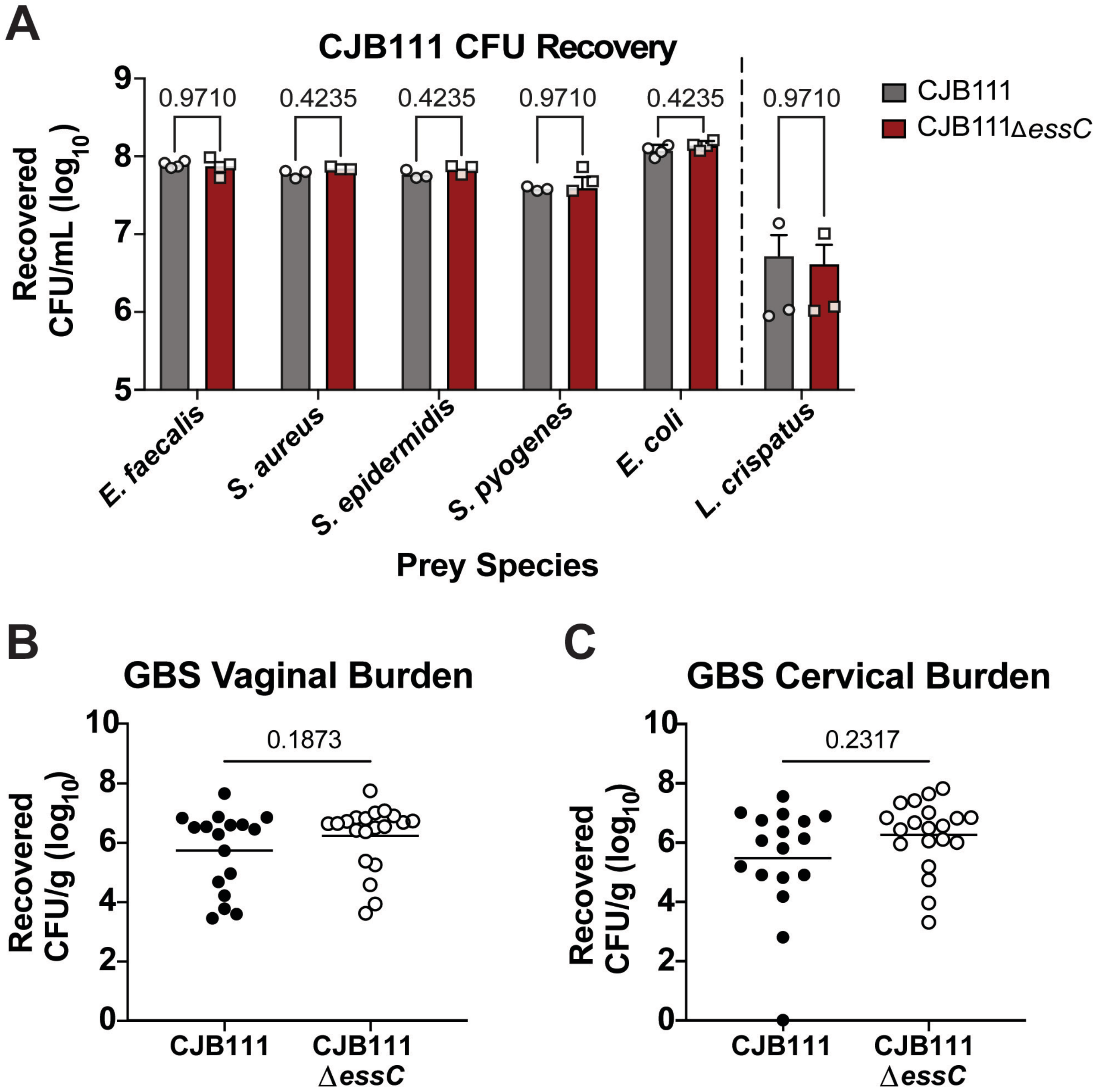

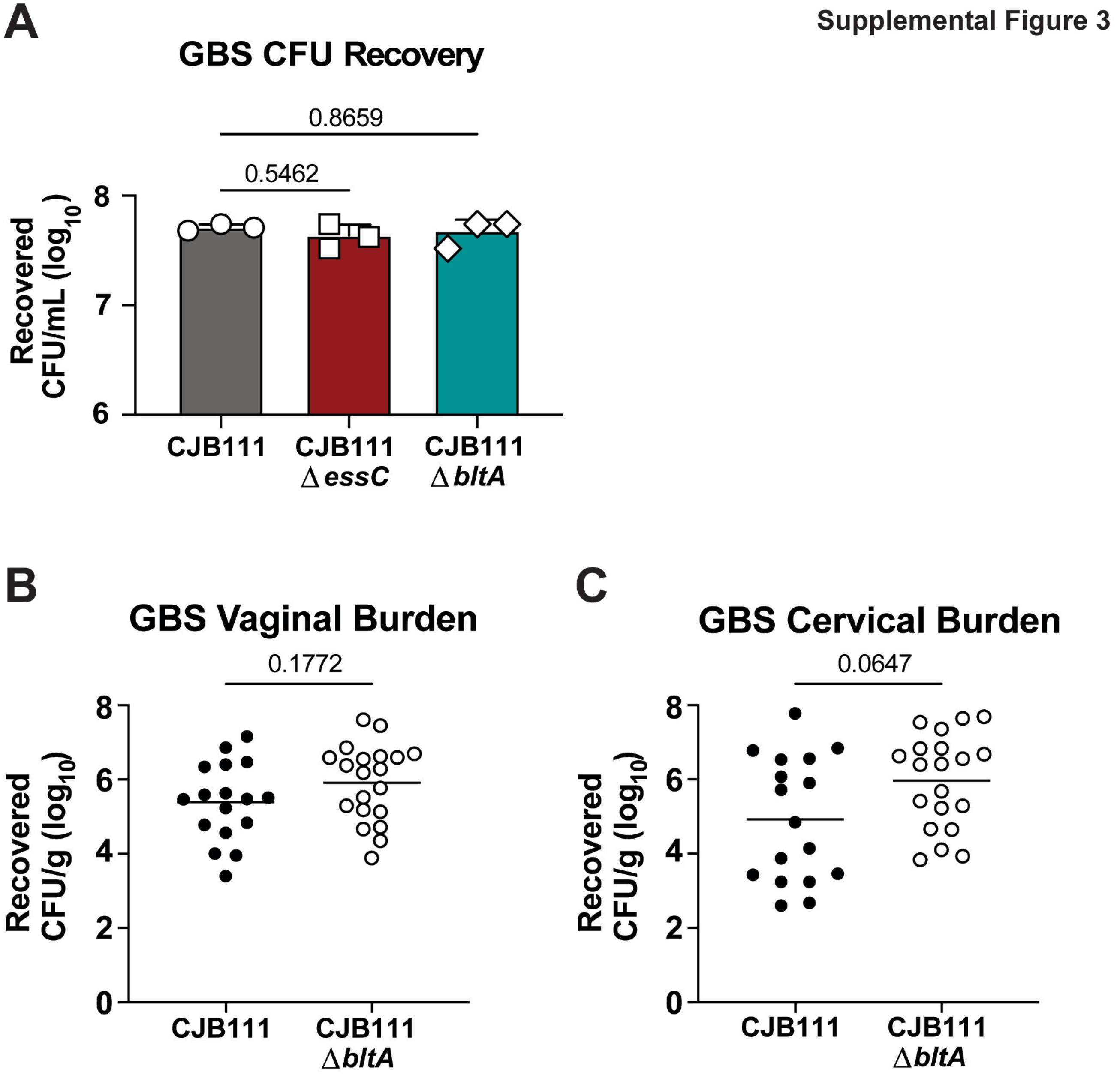

Supplemental Figure 4

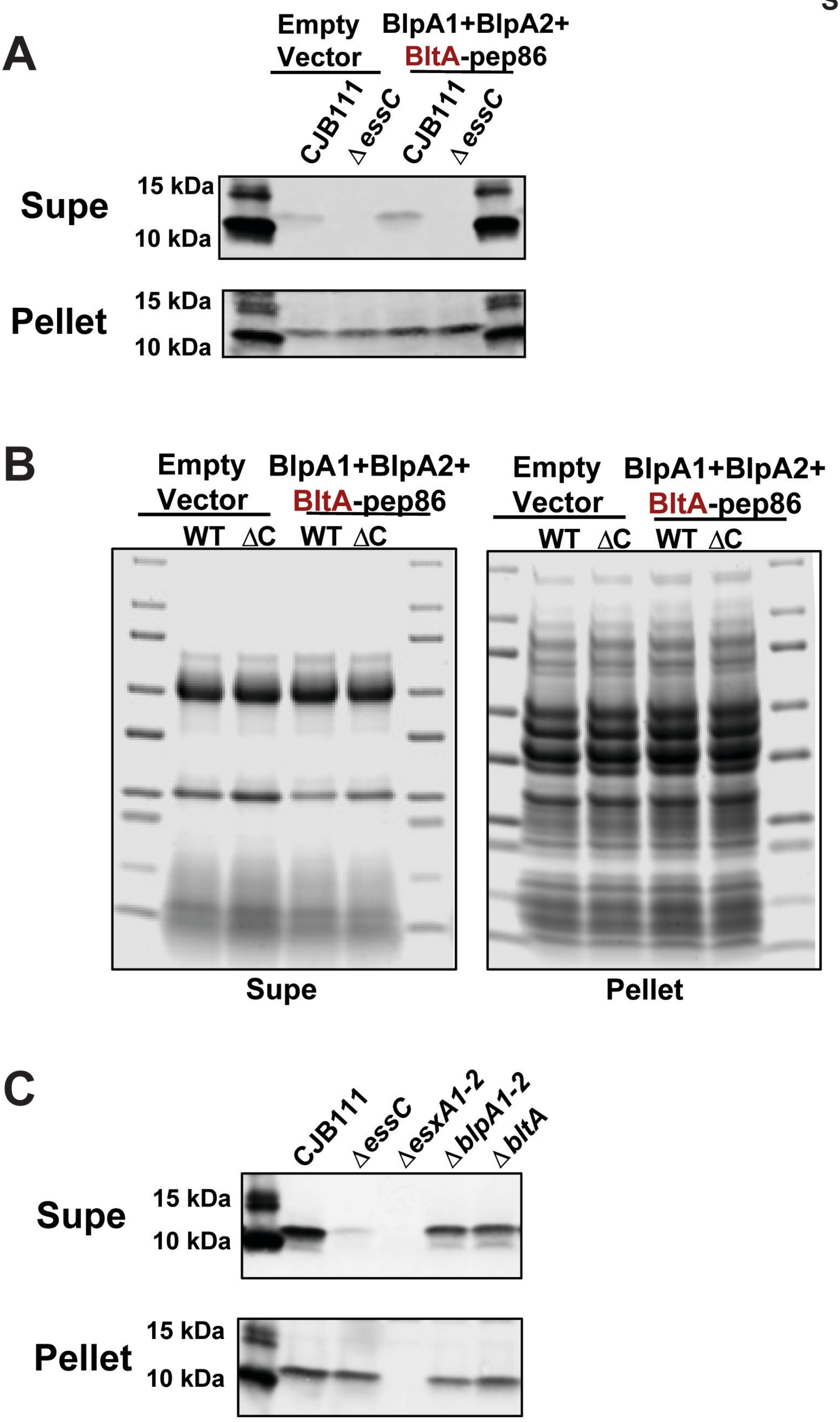

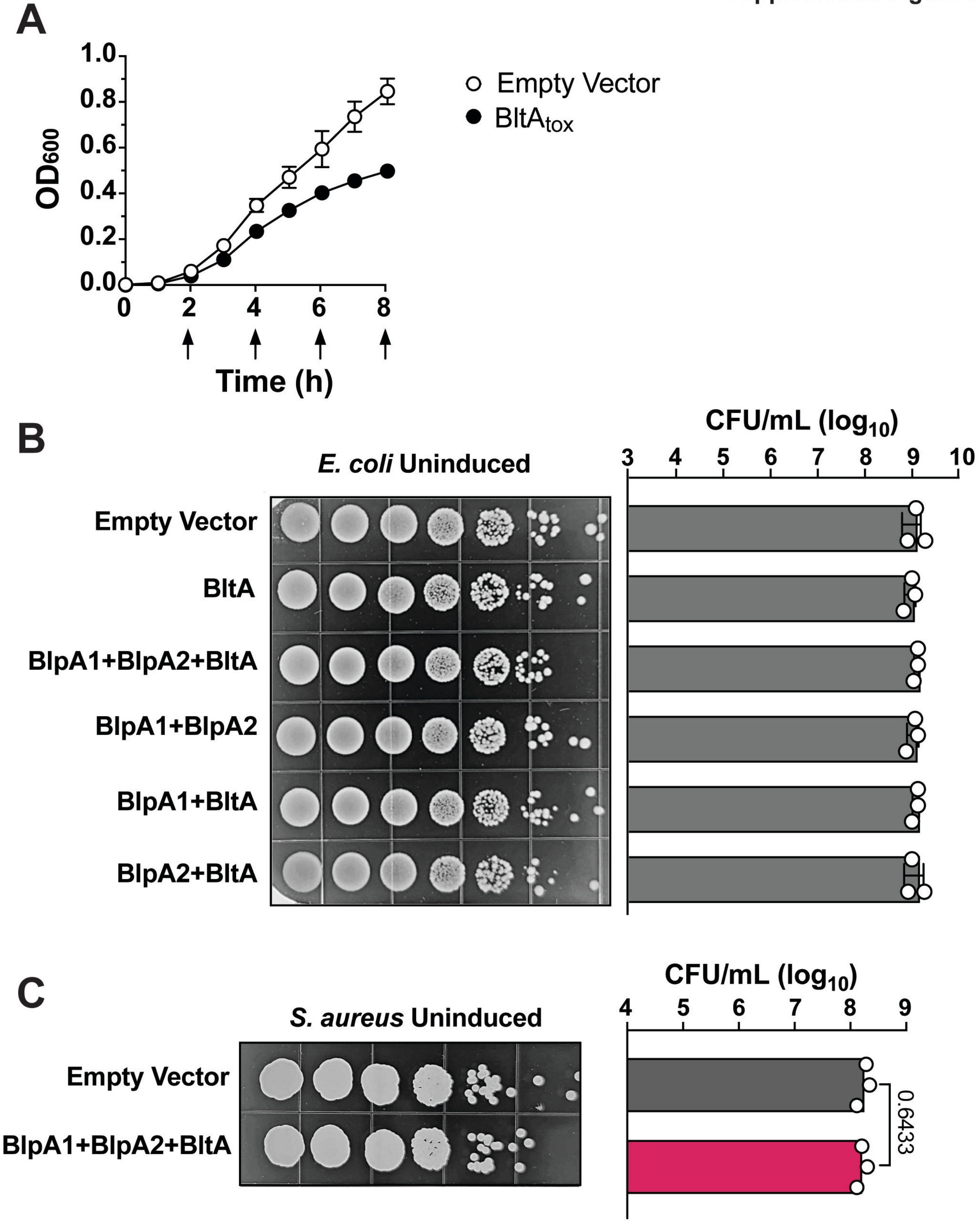

Supplemental Figure 6

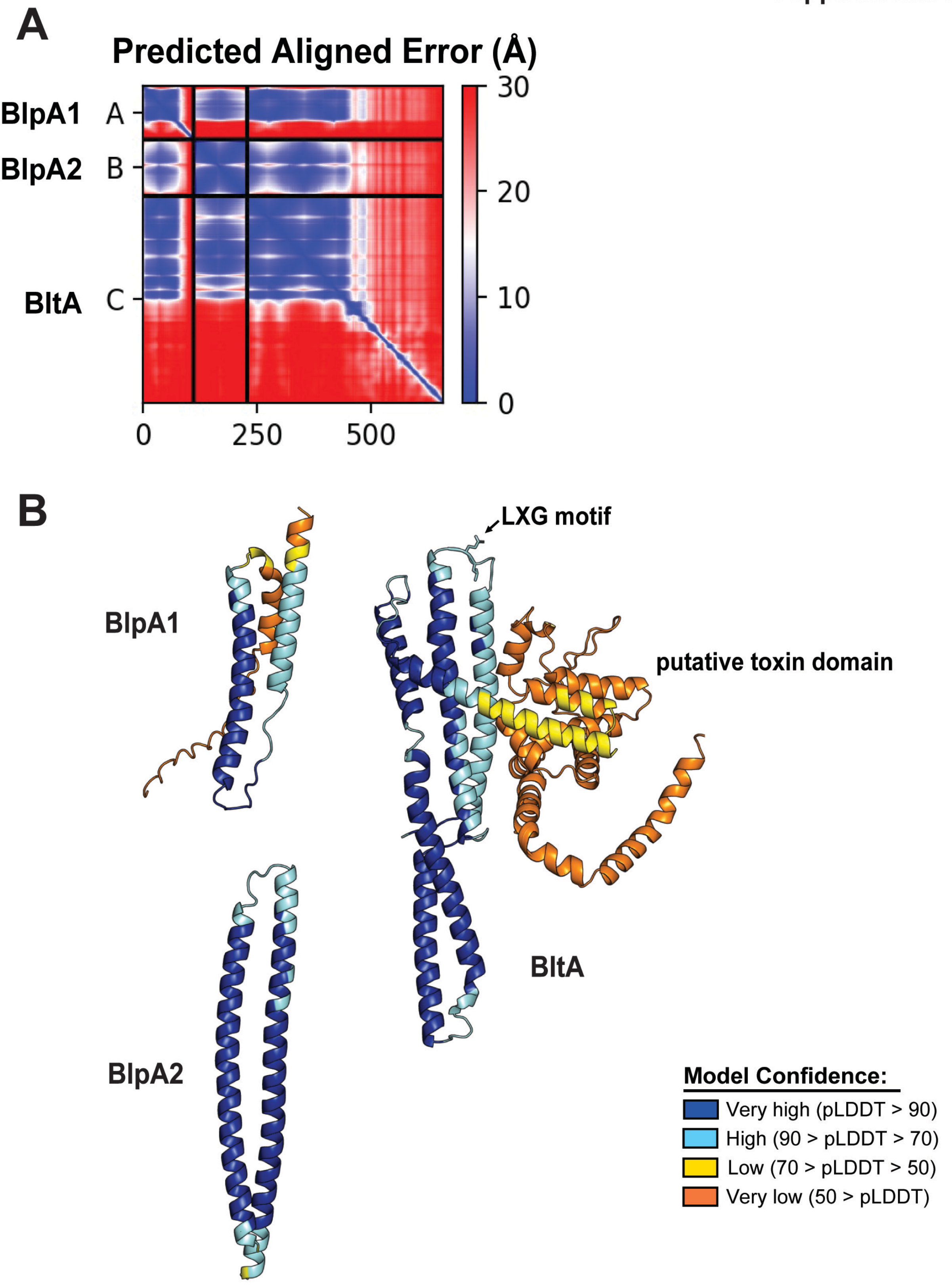

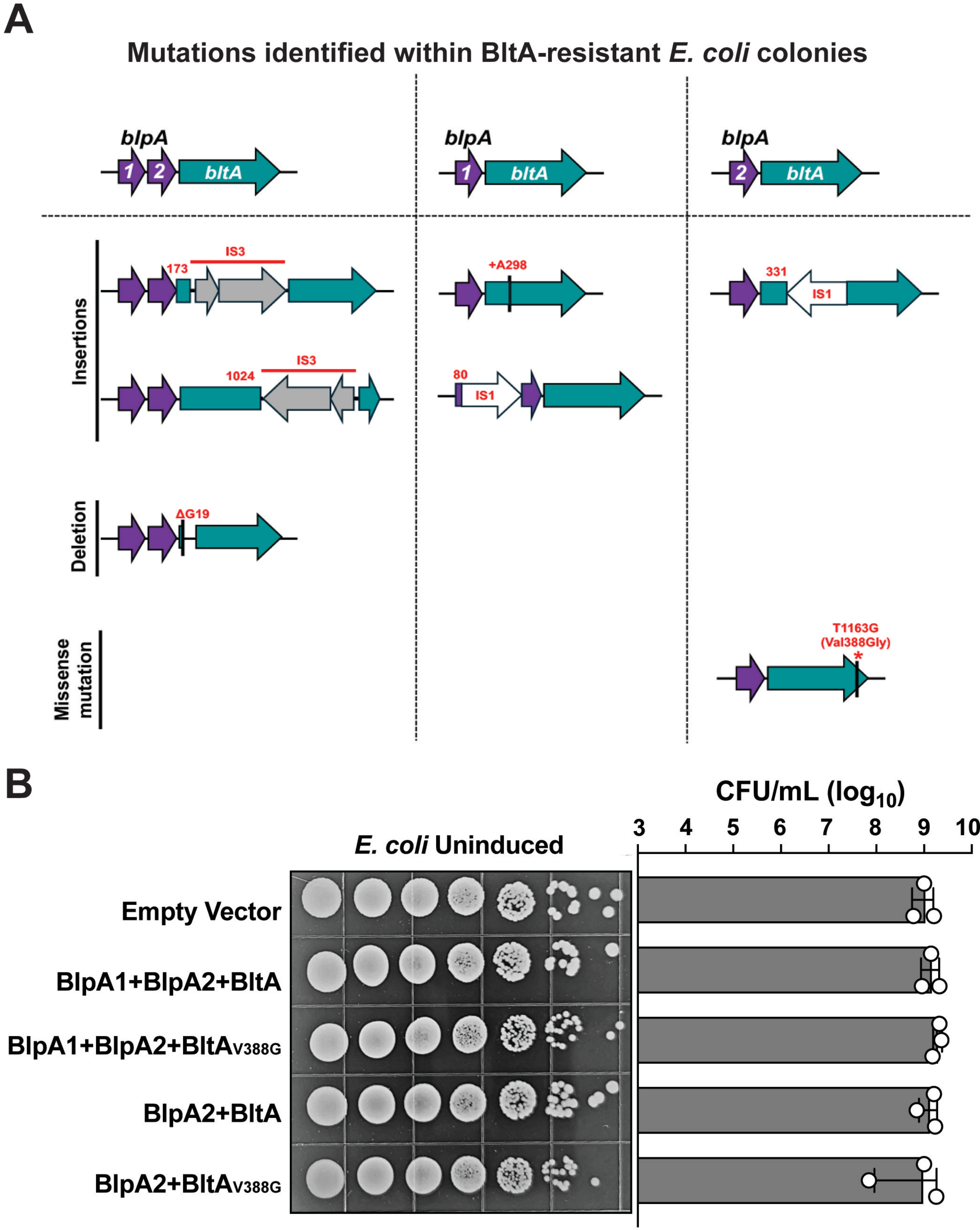
